# ERK inhibits Cic repressor function via multisite phosphorylation

**DOI:** 10.1101/2024.05.14.594119

**Authors:** Sayantanee Paul, Khandan Ilkhani, Nathan Strozewski, Liu Yang, David W. Denberg, Wootchelmine Christalin, Stanislav Y. Shvartsman, Alexey Veraksa

## Abstract

The receptor tyrosine kinase (RTK)/Extracellular Signal-Regulated Kinase (ERK) signaling pathway controls cell proliferation, differentiation, and survival. How ERK activation is relayed to its phosphorylation targets is not well understood. The transcriptional repressor Capicua (Cic) has emerged as a key target for ERK-mediated downregulation in *Drosophila* and mammals, and mutations in human *CIC* result in cancer and neurological diseases. Phosphorylation by ERK is critical for Cic downregulation, but the identities of phosphosites in *Drosophila* Cic are unknown. Here, we identify sites of phosphorylation in Cic that are directly targeted by ERK and validate their developmental functions in vivo using mutant Cic variants. Cic phosphosites are distributed throughout the length of the protein, and a group of centrally located sites appears to have a primary role in Cic downregulation. Cic mutated in 20 high-confidence sites behaves as a “super-repressor” in vivo that is largely insensitive to ERK-mediated downregulation, despite fully retaining the ability to bind to ERK. No single site is sufficient to turn off Cic activity; instead, we find that ERK must phosphorylate multiple sites in Cic simultaneously to achieve full downregulation. This multisite phosphorylation likely targets phosphodegrons that are recognized by ubiquitin ligases such as Ago/FBXW7 and contributes to Cic degradation. This study advances our understanding of the molecular mechanisms of signal interpretation downstream of the RTK/ERK signaling network.

## Introduction

Capicua (Cic) is a high-mobility group (HMG) box containing transcriptional repressor that acts downstream of the RTK/ERK signaling cascade. In *Drosophila*, ERK-mediated Cic phosphorylation and downregulation are necessary for proper patterning and growth of multiple tissues during development (Jimenez et al., 2012). In humans, mutations in *CIC* have been implicated in neurodegenerative disease spinocerebellar ataxia type 1 (SCA1) (Lam et al., 2006; Fryer et al., 2011), in the majority of oligodendroglioma cases, and other cancers (Simon-Carrasco et al., 2017; Tanaka et al., 2017; Kim et al., 2020). In both flies and mammals, Cic phosphorylation is a critical regulatory event in RTK/ERK signal transduction. Previous studies identified the C2 domain in *Drosophila* Cic that mediates its binding to ERK (Astigarraga et al., 2007), and this interaction is required for downregulation of its function as a transcriptional repressor. Several mechanisms have been proposed to explain Cic downregulation, including loss of DNA binding, export to cytoplasm, protein degradation, and loss of binding to corepressors (Astigarraga et al., 2007; Tseng et al., 2007; Ajuria et al., 2011; Dissanayake et al., 2011; Jimenez et al., 2012; Bunda et al., 2019; Keenan et al., 2020).

Since all the proposed modes of Cic inactivation are dependent on post-translational events such as ERK-dependent phosphorylation, it is essential to determine the role of these phosphosites in the context of Cic downregulation. Previous studies have identified and validated several sites of phosphorylation in human CIC, but the majority are not directly targeted by ERK. Phosphorylation of S173 near the HMG box is targeted by the kinase p90RSK downstream of ERK activation and is necessary for establishing interactions with 14-3-3 proteins, which may interfere with DNA binding (Dissanayake et al., 2011) as well as contribute to the export of CIC to the cytosol (Ren et al., 2020). A homologous site in *Drosophila* Cic (S461) was proposed to serve a similar function (Dissanayake et al., 2011). Additionally, phosphorylated S173 interacts with the ubiquitin ligase PJA1, leading to CIC degradation (Bunda et al., 2019). ERK-targeted phosphorylation of two other residues may prevent binding of a carboxy-terminal nuclear localization signal to importin, which interferes with CIC nuclear localization (Dissanayake et al., 2011). CIC is also phosphorylated by c-Src on tyrosines, which promotes its nuclear export (Bunda et al., 2020).

Despite these data, there remains a dearth of evidence regarding sites in Cic that are directly targeted by ERK, and the identities of phosphosites in *Drosophila* Cic are unknown. Here, we carried out a mass spectrometry-based screen to identify the direct ERK phosphosites in *Drosophila* Cic. We have found that Cic is phosphorylated on multiple sites and validated their functional role in vivo using variants carrying combinations of the mutated phosphosite residues. Mutation of all 20 high-confidence sites generated a “super-repressor” variant that was essentially insensitive to ERK downregulation, whereas subsets of these sites gave only a partial resistance to ERK. This suggests that Cic is regulated by ERK via multisite phosphorylation, and many sites must be targeted for complete downregulation. We propose that at least some of these ERK-dependent sites form phosphodegrons recognized by ubiquitin ligases, which may ultimately lead to Cic degradation via the proteasome.

## Results and Discussion

### Identification of ERK-dependent sites of phosphorylation in Cic

To identify the sites in Cic that are directly phosphorylated by ERK, we performed an in vitro kinase reaction using full length Cic protein tagged with streptavidin-binding peptide (SBP) (Yang et al., 2016). Cic-SBP was purified from stably transfected *Drosophila* S2 cells under the condition of MEK inhibition by PD0325901 (Ciuffreda et al., 2009) (Fig. 1A). MEK inhibition was used to reduce any background phosphorylation by endogenous ERK. Cic-SBP was immobilized on streptavidin beads and subjected to a kinase reaction using bacterially purified activated dually phosphorylated rat ERK2 (dpERK) and ATP. As a control, Cic-SBP was incubated with dpERK in the absence of ATP. The control and experimental samples were washed, eluted, and analyzed by mass spectrometry (Fig. 1A). This experiment was performed for a total of 4 biological replicates (Table S1).

**Fig. 1.**
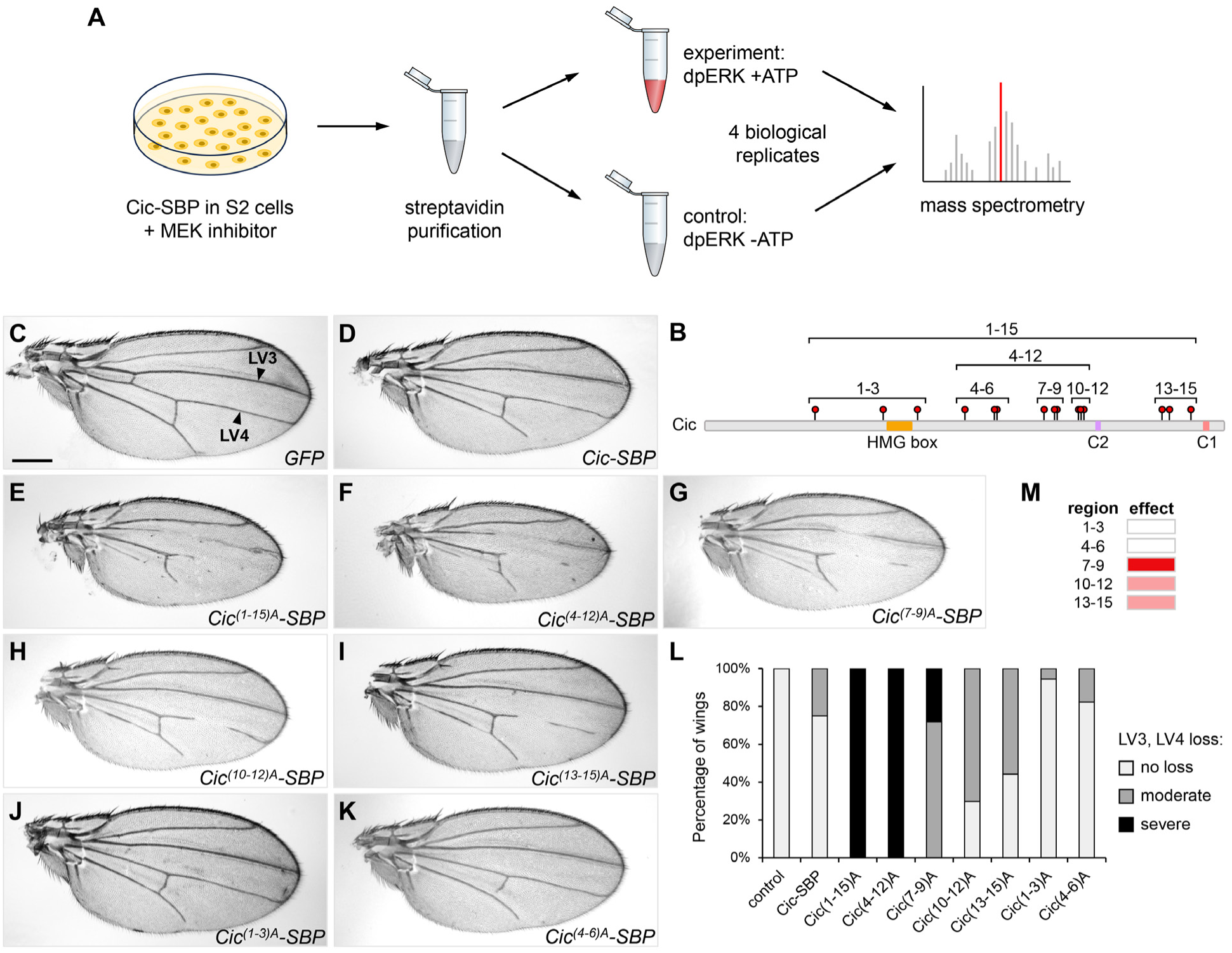
ERK-mediated multisite phosphorylation is required for Cic downregulation. (A) Experimental outline for identification of ERK-mediated phosphorylation sites in Cic. (B) Schematic showing locations and groups of mutated residues in the corresponding *UAS-Cic-SBP* constructs, based on the first mass spectrometry run (15 sites total). Individual sites are shown in Fig. S1. (C-K) Adult male wing phenotypes resulting from expression of the indicated *Cic-SBP* variants under the control of the wing-specific *MS1096-GAL4* driver. Scale bar, 300 µm. Arrowheads in (C) point to longitudinal veins 3 and 4 (LV3, LV4). (L) Quantification of LV3 and LV4 loss in (C-K). n≥27 for each genotype. Moderate loss: one incomplete and one complete LV3/4 (e.g. panel H), severe loss: both LV3 and LV4 incomplete (e.g. panel E). (M) Interpretation of the wing phenotypes in (G-K) showing relative gains over wild type Cic after mutating the indicated groups of residues, with darker color indicating strongest effect (color intensity is arbitrary).

### ERK-mediated multisite phosphorylation is required for Cic downregulation

The first round of phosphosite identification yielded 15 high confidence ERK-dependent phosphosites (serines and threonines) that were present in the experimental sample but not in the control (Fig. 1B; Fig. S1). We used this set to dissect the contributions of individual sites to Cic downregulation. Fly lines were generated that carried *UAS-Cic-SBP* with substitutions of all 15 sites to non-phosphorylatable alanines (the Cic^(1-15)A^ mutant) as well as constructs carrying subsets of these substitutions: 5 groups of 3 sites each (Cic^(1-3)A^, Cic^(4-6)A^, Cic^(7-9)A^, Cic^(10-12)A^, and Cic^(13-15)A^) and a larger set containing the three central subsets (Cic^(4-12)A^, Fig. 1B). We then asked how ERK signaling readout is affected in vivo by overexpressing these Cic phosphomutant variants in the developing wing. Since EGFR signaling relieves Cic-mediated downregulation of downstream target genes and thus promotes wing vein development (Roch et al., 2002), we focused on the loss of wing venation pattern in the adult wing as an indicator of gain of function of Cic repressor activity. Overexpression of wild type Cic using the wing pouch *MS1096-GAL4* driver (Capdevila and Guerrero, 1994) resulted in a moderate vein loss corresponding to Cic gain-of-function effect (Fig. 1C,D,L), but expression of either the Cic^(1-15)A^ or the Cic^(4-12)A^ mutant led to a much more severe phenotype (Fig. 1E,F,L), suggesting that these two mutant proteins became resistant to downregulation by ERK. Among the triplet subsets, the Cic^(7-9)A^ mutant had the strongest effect (Fig. 1G,L), followed by more C-terminally located Cic^(10-12)A^ and Cic^(13-15)A^ (Fig. 1H,I,L). Expression of Cic^(1-3)A^ and Cic^(4-6)A^ gave phenotypes that were not different from expression of wild type Cic (Fig. 1J-L).

The overall effects of the triplet subsets are summarized in Fig. 1M. While the Cic^(7- 9)A^ mutant exhibited the most severe vein loss out of the triplet combinations, it could not phenocopy the overall Cic^(1-15)A^ or the Cic^(4-12)A^ phenotype. Therefore, it appears that multiple phosphorylation sites function together to mediate ERK-dependent downregulation of Cic repressor activity downstream of EGFR.

### Cic activity in vivo correlates with the degree of phosphosite substitution

The Cic-SBP in vitro kinase reaction was repeated 3 more times, identifying 21 high-confidence phosphosites that were present in 2 or more of the 4 biological replicates (Fig. 2A; Fig. S1; Fig. S2). Based on these data, we generated a Cic^20A^ mutant variant that contained 13 sites previously included in Cic^(1-15)A^ as well as 7 sites that were not mutated in Cic^(1-15)A^ (Fig. S1). Phosphorylation of T1059 was identified twice but was omitted from mutagenesis because this site is located within the ERK binding domain, C2. Mutation of this residue impaired the binding between dpERK and Cic (Astigarraga et al., 2007), so it was likely to generate a very strong mutant in combination with other sites and obscure the effects of the other phosphosite mutations. S461, which is targeted by p90RSK/S6kII (Dissanayake et al., 2011), was not phosphorylated in our data.

**Fig. 2.**
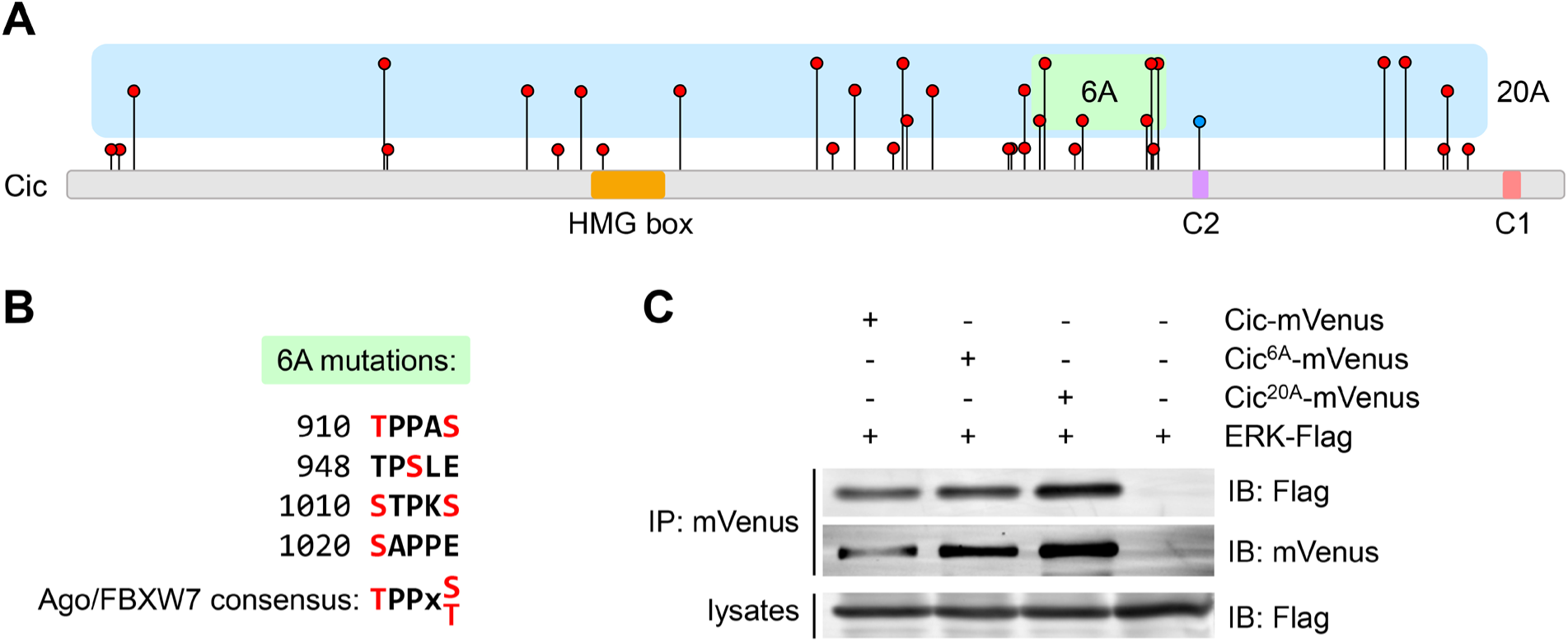
Characterization of the Cic^6A^ and Cic^20A^ mutants. (A) Schematic showing locations and identification frequency of ERK-mediated phosphorylation sites in Cic, based on 4 biological replicates. The height of site markers is proportional to the identification frequency (between 1 and 4 times). Green and blue shading shows sites chosen for mutagenesis in the Cic^6A^ and Cic^20A^ constructs, respectively. Individual sites are shown in Figs. S1 and S2. (B) Predicted Ago/FBXW7 phosphodegrons in Cic. Phosphorylated residues are shown in red. (C) Western blot showing co-immunoprecipitation between mVenus-tagged Cic variants and Flag-tagged ERK in *Drosophila* S2 cells.

Based on the in vivo results using the triplet subsets from Cic^(1-15)A^, the central region of Cic appeared to contain the sites most relevant for Cic downregulation. Phosphorylated serine and threonine residues are often encountered in phosphodegrons recognized by various E3 ubiquitin ligases (Welcker and Clurman, 2008; Holt, 2012). A previous study identified a motif in the middle portion of Cic that conforms to the consensus recognition sequence (pTPPxpS/T) of the E3 ubiquitin ligase Archipelago (Ago), which is homologous to mammalian FBXW7/Cdc4 (Suisse et al., 2017; Singh et al., 2022). We identified three additional regions nearby that generally conform to this consensus sequence and made a corresponding construct (named Cic^6A^) that specifically targeted the phosphosites included in these four putative phosphodegrons (Fig. 2B; Fig. S2).

We then thought to verify that the mutation of these phosphosites did not disrupt the binding between Cic and ERK, as the observed effects could be due simply to a loss of this interaction. To test this, we co-expressed mVenus-tagged wild type and mutant Cic constructs with *Drosophila* ERK-Flag in S2 cells and assayed their binding by co-immunoprecipitation. As shown in Fig. 2C, mutations of the phosphosites included in these constructs did not affect the binding of Cic to ERK, suggesting that any phenotypes resulting from overexpression of these constructs would not stem from an inability of ERK to associate with these Cic variants.

Overexpression of wild type Cic-mVenus in the wing using *MS1096-GAL4* resulted in a mild vein loss corresponding to Cic gain-of-function effect (Fig. 3A,B), but expression of either the Cic^6A^ or Cic^20A^ mutant led to a much more severe loss of veins, suggesting that these two mutant proteins are resistant to downregulation by ERK (Fig. 3C,D). Strikingly, Cic^20A^ had a phenotype that was almost as strong as the one caused by the Cic^ΔC2^ mutant that abrogates ERK interaction (Fig. 3D,E). A similar trend was observed when the Cic^6A^ and Cic^20A^ phosphomutant variants were expressed in the eye, where Cic is also downregulated by EGFR/ERK signaling during development (Tseng et al., 2007). Expression of Cic variants with progressive increase of phosphosite substitutions using the eye-specific driver *GMR-GAL4* led to a corresponding increase in the severity of eye loss (Fig. 3F-J, quantified in Fig. 3K,L). As in the wing, the Cic^ΔC2^ mutant gave the strongest phenotype in the eye (Fig. 3J,L).

**Fig. 3.**
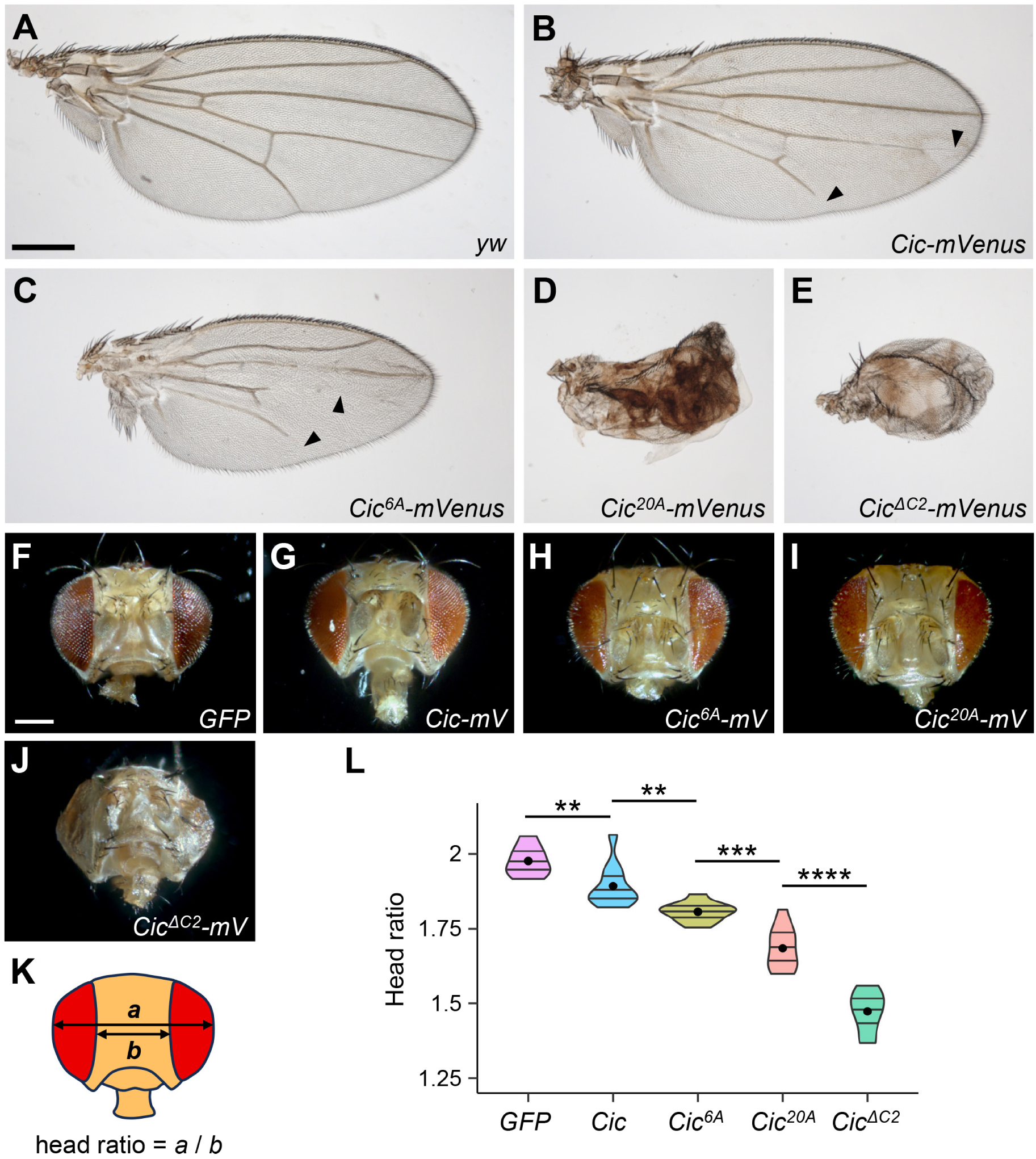
Cic activity in vivo correlates with the degree of phosphosite substitution. (A-E) Adult female wing phenotypes resulting from expressing the indicated *Cic-mVenus* variants under the control of the wing-specific *MS1096-GAL4* driver. Arrowheads indicate vein loss. Scale bar in (A), 300 µm. (F-J) Adult female head phenotypes resulting from expressing the indicated *Cic-mVenus* (*mV*) variants using the eye-specific *GMR-GAL4* driver. Scale bar in (F), 200 µm. (K, L) Quantification of the head phenotypes shown in (F-J). n≥10 for each genotype. Statistics calculated by ANOVA (*F* = 97.81, *p* < 0.001) followed by post-hoc Student’s t-tests (\*\**p* < 0.01, \*\*\**p* < 0.001, \*\*\*\**p* < 0.0001) using ggpubr and ggplot2 in RStudio.

Collectively, these results suggest that the Cic^20A^ mutations eliminated most of the phosphosites important for Cic downregulation by ERK. At the same time, the Cic^6A^ mutations did not account for all of the effects of Cic phosphorylation by ERK, and additional sites included in Cic^20A^ strongly contributed to downregulation. Since the Cic^ΔC2^ mutant exhibited a further increase in phenotype severity, there may be additional functional phosphosites present in Cic (e.g. those that were identified once in our mass spectrometry analysis) that were not included in Cic^20A^.

### ERK-mediated multisite phosphorylation of Cic is required for Cic degradation and target gene expression

Downregulation of Cic by ERK is well characterized in the early embryo, where spatially restricted activation of Torso induces phosphorylation and activation of ERK, leading to a reduction of Cic levels at both poles of the embryo, presumably through proteolytic degradation (Astigarraga et al., 2007; Grimm et al., 2012). We generated transgenic fly lines carrying *pTIGER*-based mVenus-tagged Cic phosphomutants and expressed them using the maternal driver *MTD-GAL4* to study their effects in the early embryo. The wild-type Cic-mVenus was properly downregulated in the cell nuclei at the poles where Torso is active, compared to the higher nuclear levels observed in the middle of the embryo (Fig. 4A-A’’). The Cic^6A^ and the Cic^20A^ mutants exhibited prominent nuclear signal at the poles (Fig. 4B-C’’) suggesting that these phosphomutants are less sensitive to downregulation by ERK. In this assay, the phenotype for Cic^20A^ was indistinguishable from that of Cic^ΔC2^ (Fig. 4C-D’’).

**Fig. 4.**
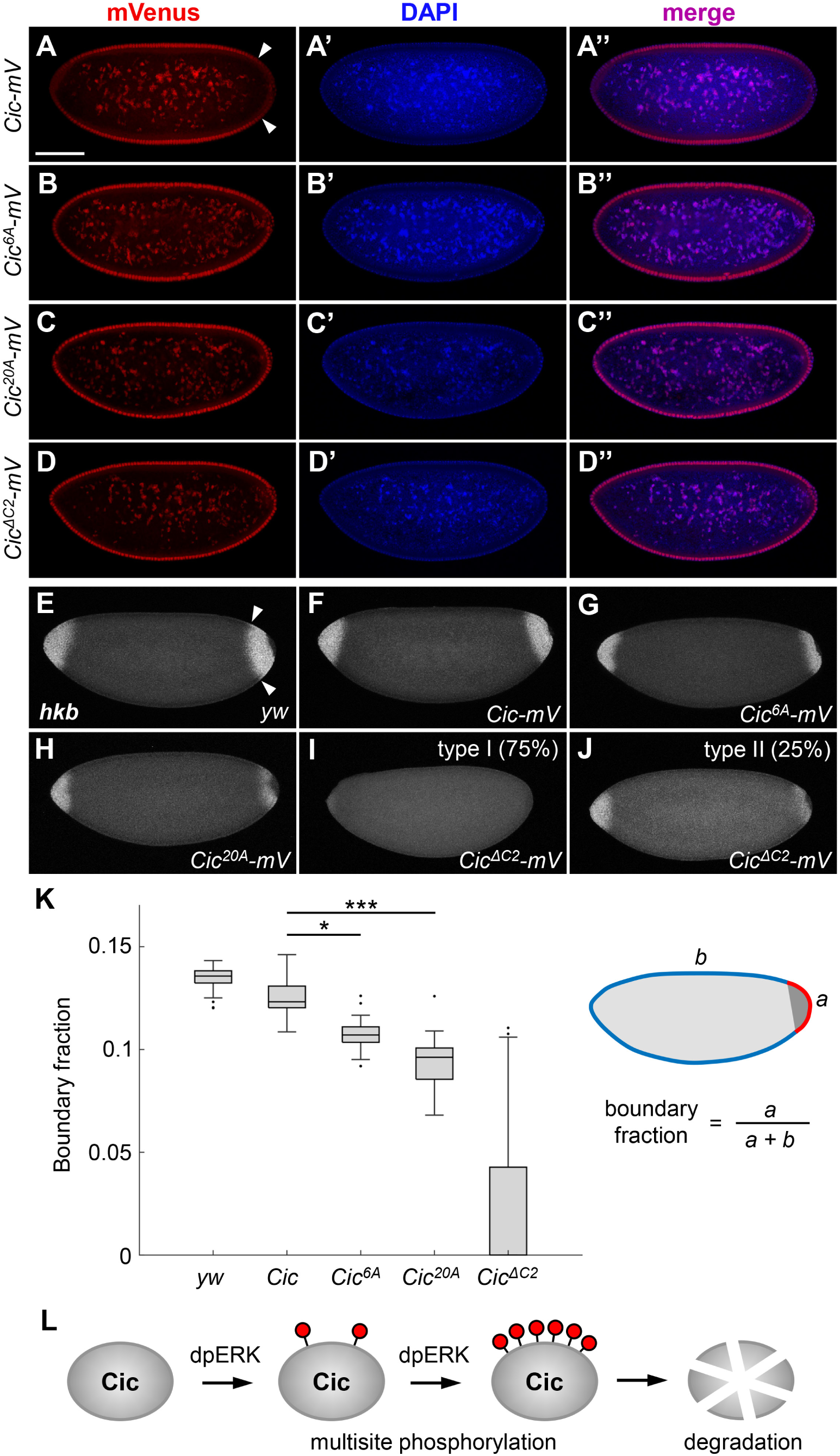
ERK-mediated multisite phosphorylation of Cic is required for Cic degradation and target gene expression. (A-D’’) Cic-mVenus immunofluorescence in embryos collected from females expressing the indicated *Cic-mVenus* variants under the control of the maternal MTD-GAL4 driver. Red, mVenus signal; blue, DAPI (DNA) signal. Scale bar in (A), 100 µm. Arrowheads in (A) delimit the posterior domain of Cic degradation. (E-J) Fluorescence in situ hybridization for *huckebein* (*hkb*) in nuclear cycle 14 embryos collected from females expressing the indicated *Cic-mVenus* variants under the control of the maternal *MTD-GAL4* driver. *hkb* expression was eliminated in the posterior region in 75% (21 out of 28) of embryos derived from the *MTD>Cic^ΔC2^* females (I, J). (K) Quantification of *hkb* expression shown in (E-J). n≥23 for each genotype. The length of the posterior *hkb* expression region (*a*, delimited by arrowheads in panel E) was measured relative to the embryo perimeter (*a+b*) and represented as boundary fraction. The embryo was segmented using an active contour technique, and the *hkb* expressing region was identified along the boundary contour by its change in intensity relative to non-expressing regions of *hkb* (see Materials and Methods). A multiple comparison Tukey-Kramer test was used for statistical analysis. \**p* < 0.05, \*\*\**p* < 0.001. Error bars denote standard error of the mean. (L) Summary model. ERK-mediated multisite phosphorylation of Cic results in downregulation of Cic activity, at least in part via degradation.

At the embryo poles, downregulation of Cic allows for expression of downstream Torso pathway target genes such as *huckebein* (*hkb*) (Jimenez et al., 2000). We studied the expression pattern of *hkb* in embryos with maternally provided expression of Cic phosphomutants using fluorescent in situ hybridization. Wild type Cic-mVenus slightly decreased the posterior domain of *hkb* expression, and expression of the Cic^6A^ or Cic^20A^ mutants showed a significant further decrease (Fig. 4E-H,K). Expression of Cic^ΔC2^ resulted in a severe loss of *hkb* expression, with 75% of embryos lacking the signal altogether (Fig. 4I-K). These data show that the phosphosites we identified are required for ERK-mediated downregulation and relief of repression that allows target gene expression in the Torso/ERK signaling pathway. As in the other assays, *hkb* expression analysis demonstrated that multisite phosphorylation of Cic by ERK is critical for achieving the proper level of downregulation.

## Conclusion

In this work, we have identified phosphosites in Cic that are directly targeted by ERK and tested their functions in vivo via expression of mutagenized Cic variants. These assays involve developmental processes that are dependent on ERK-mediated downregulation of Cic activity as a transcriptional repressor, such as the EGFR-dependent patterning of the wing veins and the Torso-dependent gene expression in the embryo. Consistently, across these assays, the Cic^6A^, Cic^20A^, and Cic^(1-15)A^ variants behaved as gain of function Cic mutants with a stronger repressor activity than wild type Cic (“super-repressors”), indicated by the mutant phenotypes. Cic^ΔC2^ was the strongest repressor because it impairs binding to ERK. Importantly, partial sets of phosphosite mutations gave weaker phenotypes, indicating the requirement for multisite phosphorylation for proper Cic downregulation.

We propose that such multisite phosphorylation is necessary for Cic degradation (Fig. 4L), as suggested by the embryo data. The *Drosophila* FBXW7/Cdc4 ortholog Archipelago (Ago) is required for targeting Cic for degradation in flies (Suisse et al., 2017). The sites that we mutated within the Cic^6A^ and Cic^(7-9)A^ subsets are similar to the FBXW7 phosphodegron consensus sequence pTPPxpS/T, where the first pT is the critical phosphorylated residue, and the second pS/T, also occasionally represented by the negatively charged D or E, helps in recognition (Singh et al., 2022). These regions in Cic appear to be dominant in mediating ERK-dependent downregulation (Fig. 1M), however other phosphosites are required for the full effect.

Phosphodegron-mediated degradation has been shown to be an important mechanism for the regulation of transcriptional activators and repressors in both yeast and mammalian systems (Liu et al., 2010; Holt, 2012). The homology between human and fly Cic orthologs is poor outside the conserved HMG box and C1 domains, making it difficult to align the phosphosites in *Drosophila* Cic with the human CIC sequence. Instead, we have identified six possible phosphodegrons in human CIC that are located in the carboxy-terminal half of the protein, in a similar general location as the phosphodegrons in fly Cic (Fig. S3). Some of the serine and threonine residues in these motifs were found to be phosphorylated in high-throughput studies (Fig. S3, PhosphoSitePlus). We speculate that human CIC downregulation may occur at least in part via these sequences.

A recent study proposed that multisite phosphorylation may result in intramolecular repulsion in mouse CIC, which in turn would lead to its dissociation from the DNA and promote cytoplasmic localization (Park et al., 2022). Our data suggest that in addition to these effects, multisite phosphorylation may also contribute to mammalian CIC degradation.

## Materials and Methods

### Drosophila melanogaster stocks

Fly stocks and crosses were maintained on standard yeast-cornmeal-agar medium at 25°C or 18 °C. The following driver lines were used: *MS1096-GAL4* (wing pouch) (Capdevila and Guerrero, 1994), *GMR-GAL4* (eye) (Hay et al., 1997), *MTD-GAL4* (maternal triple driver) (Mazzalupo and Cooley, 2006). Transgenic lines were generated by inserting the constructs into the *attP2* genomic site using the φC31-based integration system (Venken et al., 2006; Bischof et al., 2007). Injections were performed by Rainbow Transgenic Flies, Inc. All constructs were integrated in the same site and therefore provide a matching set for comparisons of expression levels and phenotypes.

### Plasmid construction

Construction of carboxy-terminally tagged full length *pMK33-Cic-SBP* and *pUAST-Cic- SBP* was described in (Yang et al., 2016). For making Cic^(1-15)A^ and Cic^20A^, gene fragments with mutations were synthesized by Twist Biosciences. Construction of *pUAST-attB-Cic^(1-15)A^-SBP* and *pUAST-attB-Cic^20A^-mVenus* was carried out by assembling gene fragments using the HiFi DNA assembly Kit (NEB). Construction of the other subgroups of Cic mutants was done by assembling different parts of Cic generated by overlap PCR using wild type Cic, Cic^(1-15)A^, and Cic^20A^ as templates. For maternal expression, Cic variants were cloned into pTIGER vector (Ferguson et al., 2012). Construction of carboxy-terminally tagged *ERK-Flag* was described in (Tipping et al., 2010). For co-immunoprecipitation experiments, wild type Cic and mutant variants were subcloned into pMT-V5-His vector (Invitrogen).

### *In vitro* kinase reactions

For stable expression in S2 cells, pMK33-Cic-SBP construct was transfected by using Effectene transfection reagent (Qiagen), and stable cell lines were selected in the presence of 300 μg/mL hygromycin (Sigma), as described in (Yang and Veraksa, 2017). *pMK33-Cic-SBP* stable cells were pre-incubated with 2 μM PD0325901, a MEK inhibitor (Biotang Inc.), with DMSO as vehicle, for 3 hrs before induction. Cells were induced with 0.35 mM CuSO_4_ overnight. Cells were harvested and then lysed with default lysis buffer (50 mM Tris pH 7.5, 125 mM NaCl, 5% glycerol, 0.2% IGEPAL CA-630, 1.5 mM MgCl_2_, 1 mM DTT, 25 mM NaF, 1 mM Na_3_VO_4_, 1 mM EDTA) containing 2x Complete protease inhibitor (Roche). Cleared cell lysates were incubated with Streptavidin beads (Pierce) at 4 °C for 2 hrs. After three washes (last wash in kinase buffer), 500 ng of purified dpERK and 200 mM ATP in kinase buffer (Cell Signaling Technologies) were added and samples were incubated at 30°C for 30 min. dpERK without ATP was used as a negative control. Purification of phosphorylated ERK from bacteria was described in (Paul et al., 2020). After several washes, samples were eluded with 2xSDS sample buffer, analyzed by 6% SDS-PAGE gel, and Cic phosphorylation was analyzed by nanoLC-MS/MS at the Taplin Mass Spectrometry Facility at Harvard Medical School.

### Identification of phosphorylation sites by mass spectrometry

Excised gel bands were cut into approximately 1 mm^3^ pieces. The samples were reduced with 1 mM DTT for 30 min at 60°C and alkylated with 5 mM iodoacetamide for 15 min in the dark at room temperature. Gel pieces were then subjected to a modified in-gel trypsin digestion procedure. Gel pieces were washed and dehydrated with acetonitrile for 10 min followed by removal of acetonitrile. Pieces were then completely dried in a speed-vac. Rehydration of the gel pieces was with 50 mM ammonium bicarbonate solution containing 12.5 ng/µl modified sequencing-grade trypsin (Promega, Madison, WI) at 4°C. Samples were then placed in a 37°C room overnight. Peptides were later extracted by removing the ammonium bicarbonate solution, followed by one wash with a solution containing 50% acetonitrile and 1% formic acid. The extracts were then dried in a speed-vac (∼1 hr). The samples were then stored at 4°C until analysis.

On the day of analysis the samples were reconstituted in 5-10 µl of HPLC solvent A (2.5% acetonitrile, 0.1% formic acid). A nano-scale reverse-phase HPLC capillary column was created by packing 2.6 µm C18 spherical silica beads into a fused silica capillary (100 µm inner diameter x ∼30 cm length) with a flame-drawn tip. After equilibrating the column each sample was loaded via a Famos auto sampler (LC Packings, San Francisco CA) onto the column. A gradient was formed and peptides were eluted with increasing concentrations of solvent B (97.5% acetonitrile, 0.1% formic acid).

As each peptide was eluted, they were subjected to electrospray ionization and then they entered into an LTQ Orbitrap Velos Pro ion-trap mass spectrometer (Thermo Fisher Scientific, San Jose, CA). Eluting peptides were detected, isolated, and fragmented to produce a tandem mass spectrum of specific fragment ions for each peptide. Peptide sequences (and hence protein identity) were determined by matching protein or translated nucleotide databases with the acquired fragmentation pattern by the software program, Sequest (ThermoFinnigan, San Jose, CA) (Eng et al., 1994). The modification of 79.9663 mass units to serine, threonine, and tyrosine was included in the database searches to determine phosphopeptides. Phosphorylation assignments were determined by the Ascore algorithm (Beausoleil et al., 2006). All databases include a reversed version of all the sequences and the data was filtered to between a one and two percent peptide false discovery rate.

### Co-immunoprecipitation and western blotting

Wild type and mutant Cic-mVenus variants and ERK-Flag were expressed from the pMT vector-based constructs in cultured *Drosophila* S2 cells. S2 cells were cultured at 25°C in standard Schneider’s S2 medium with 10% FBS (Gibco) and 5% Pen/Strep (Invitrogen). Proteins were induced with 0.35 mM CuSO_4_ overnight, cells were lysed in default lysis buffer as above, and protein complexes were isolated using GFP-Trap beads (Bulldog Bio). After washes and elution with 4xSDS sample buffer, protein complexes were resolved on 7% SDS protein gels and transferred onto Millipore Immobilon-FL PVDF Transfer Membranes with 0.45 μm pores. Primary antibodies used for western blots were as follows: mouse anti-Flag 1:1,000 (Sigma), rabbit anti-GFP 1:1,000 (ThermoFisher). Secondary antibodies used were as follows: IRDye 800CW Donkey anti-Rabbit IgG 1:10,000 (LI-COR), and IRDye 680CW Goat anti-Mouse IgG, 1:10,000 (LI-COR).

### Immunohistochemistry

To assess embryonic Cic-mVenus localization, 0-4 hr embryos collected from *MTD>yw*, *MTD>Cic-mVenus*, *MTD>Cic^6A^-mVenus*, *MTD>Cic^20A^-mVenus* and *MTD>Cic^ΔC2^-mVenus* mothers were dechorionated with 50% (v/v) Clorox bleach, rinsed with water, then fixed for 20 minutes at room temperature (RT) in a fixative comprised of 4% (v/v) paraformaldehyde (Electron Microscopy Sciences), 25% (v/v) Heptane, 1x PBS, and MilliQ water. After fixation, the embryos were devitellinized via the addition of 8 mL of methanol and harsh agitation for 90 seconds. Fixed and devitellinized embryos were collected, washed three times in methanol, four times in ethanol, then stored in ethanol at -20°C. Embryos were rehydrated once with ethanol, twice with 1:1 ethanol:PBT (1x PBS with 0.1% (v/v) Tween 20), then twice with 1x PBT. The embryos were incubated in blocking reagent (1:1 (v/v) Roche Blocking Reagent and 1x PBT) for 2 hrs at RT and incubated overnight at 4°C in primary antibody solution (1:100 rabbit anti-GFP (ThermoFisher) in blocking reagent). Embryos were washed at RT in 0.1% BSA (w/v in 1x PBT), blocked for 1 hr then incubated with secondary antibody solution (1:500 goat anti-rabbit Alexa Fluor 555 (ThermoFisher) in blocking reagent) for 2 hrs in the dark at RT. Embryos were then washed in the dark then mounted in Prolong Gold Antifade Mountant with DAPI (ThermoFisher). Images were acquired with Zeiss LSM 880 confocal microscope.

### *hkb* fluorescent in situ hybridization (FISH) and image analysis

0-4 hr embryos were collected as above and used for FISH experiments using standard FISH protocols. In brief, ∼50 µl of fixed embryos were incubated in 90% xylenes for 1 hr, followed by wash steps with ethanol, methanol and PBT. Embryos were incubated at 65°C with hybridization buffer (50% formamide, 5x SSC, 100 µg/ml sonicated salmon sperm DNA, 50 µg/ml heparin, 0.1% Tween 20) for 4 hrs. The samples were resuspended with Digoxigenin (DIG) labeled antisense *hkb* RNA probes with hybridization buffer (1:25) and incubated at 65 °C overnight. After hybridization, the samples were washed with hybridization buffer and PBST, followed by standard immunostaining protocols. DIG labeled antisense *hkb* RNA probe was synthesized by amplification of the *hkb* cDNA. Nuclear cycle 14 embryos were selected for imaging. DAPI was used for staining nuclei. Sheep anti-DIG (1:25; Roche) was used as primary antibody and Alexa Fluor conjugate (1:500; Invitrogen) was used as secondary antibody. Imaging for FISH experiments was performed on a Leica SP5 confocal microscope with following specifications: 20x AIR objective, 405-nm and 561-nm diode lasers.

Segmentation of the *Drosophila* embryo perimeter was performed using the following procedure. From the *hkb*-labeled image, Otsu’s method was used to approximately separate the interior and exterior of the embryo. This was followed by a flood fill operation to fill holes in the mask and a morphological dilation. This binary mask was used as input along with the raw image to an active contour technique to finetune the segmentation of the embryo’s interior. The boundary of this mask, the embryo’s perimeter, was isolated as a piecewise parametric curve which was smoothed using a Savitzky-Golay filter. We quantified *hkb* intensity at the embryo border by averaging pixel intensity values within a Euclidean distance of 20 pixels (12.61 µm) of the parametric boundary curve and in the interior of the embryo. Finally, we used MATLAB signal processing toolbox functions ‘risetime’ and ‘falltime’ to determine the position on the curve where the *hkb* pole starts and stops. These functions estimate the time instant of a state transition within a signal. Code is available at the GitHub repository: https://github.com/ddenberg/HKB-Quantification.

### Wing and head phenotypes

Adult wings and heads were imaged with Olympus BX60 compound microscope using bright-field illumination and 4x objective. For the SBP-tagged Cic^(1-15)A^ series mutants, wings from male progeny of crosses with *MS1096-GAL4* were analyzed. For the mVenus-tagged Cic^6A^, Cic^20A^, and Cic^ΔC2^ mutants, wings and heads from female progeny from the crosses with *MS1096-GAL4* or *GMR-GAL4* were analyzed.

## Acknowledgements

We thank the Bloomington *Drosophila* Stock Center for their services. Mass spectrometry was performed at the Taplin Mass Spectrometry Facility at Harvard Medical School. This work was supported by the NIH grant GM141843 to A.V. and S.Y.S. S.P. was supported by the Sanofi and Oracle UMass Boston Doctoral Fellowships.

---

**Fig. S1.**
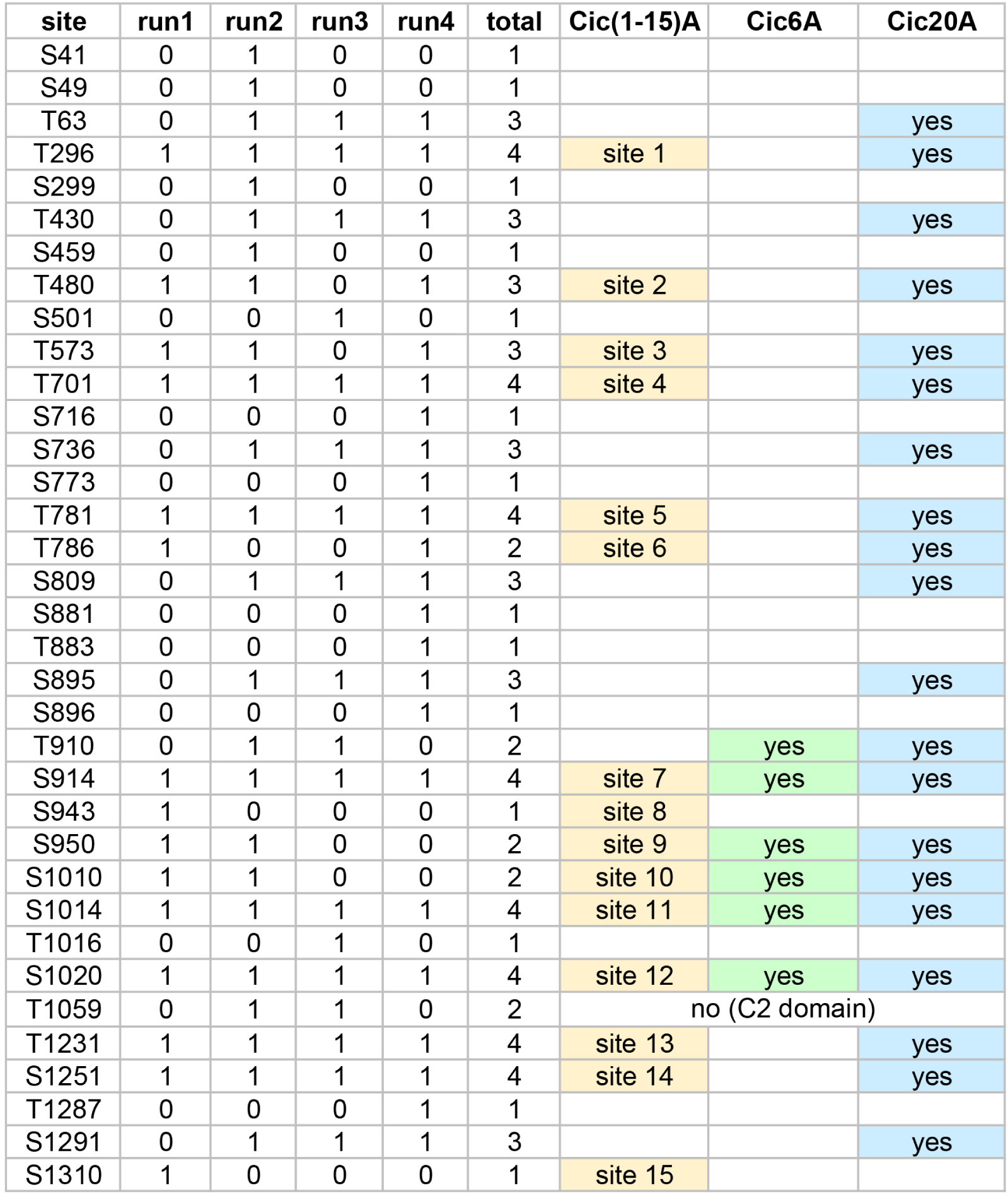
ERK-mediated Cic S/T phosphorylation sites identified by mass spectrometry (biological replicates 1-4) and sites chosen for mutagenesis. The three rightmost columns show the sites mutated to alanine in the corresponding Cic variants: Cic^(1-15)A^ was based on the first biological replicate (run 1), and Cic^6A^ and Cic^20A^ were based on all four runs. Cic^20A^ includes all sites identified in at least two runs, except T1059 located in the C2 domain, which is the binding site for ERK. Cic^6A^ is a subset of Cic^20A^. Mass spectrometry data are shown in Table S1.

**Fig. S2.**
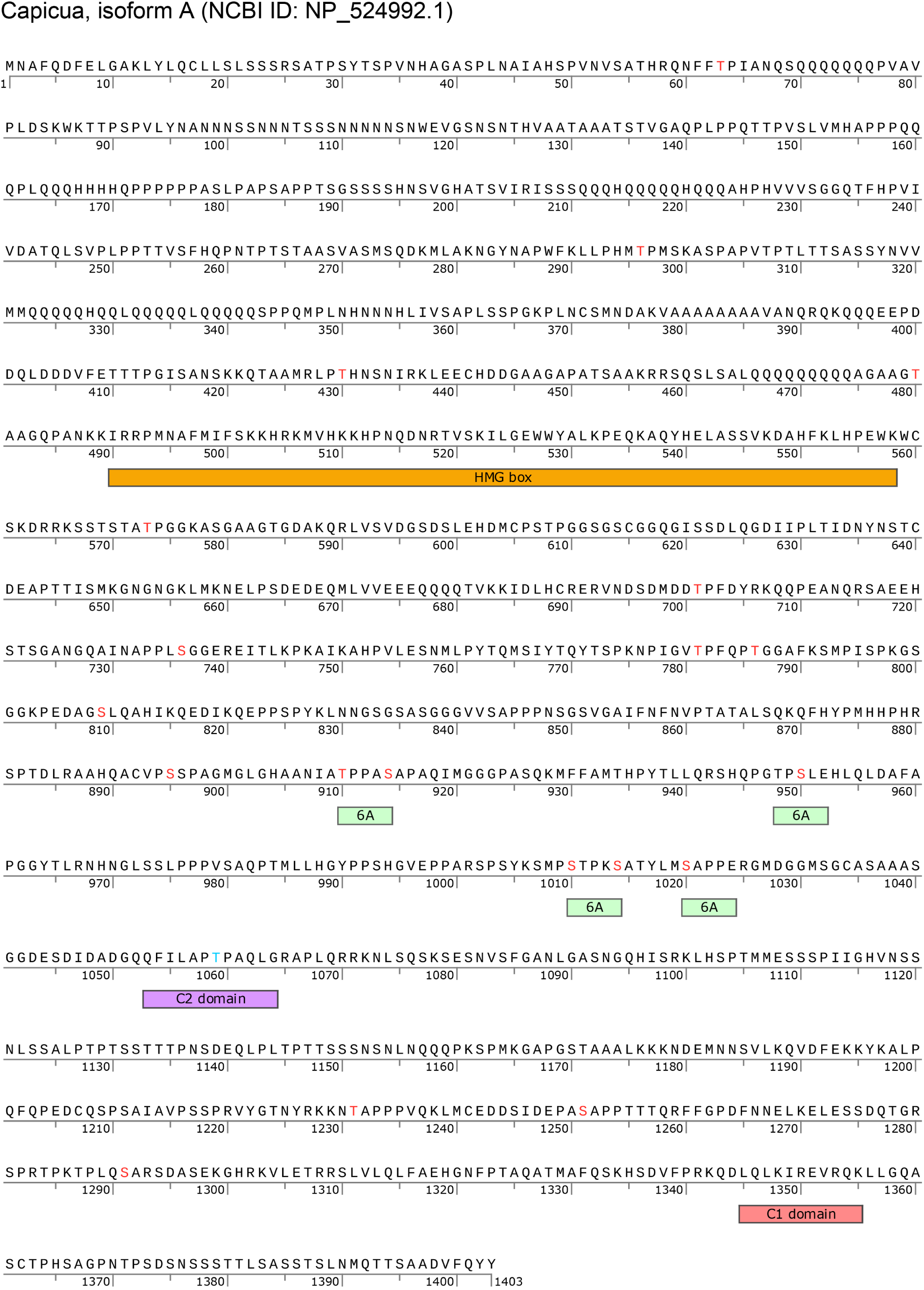
Annotated sequence of the short isoform of Capicua (isoform A) used in this study. S/T phosphorylation sites mutated in Cic^20A^ are shown in red. Green boxes labeled “6A” show putative Ago/FBXW7 phosphodegrons and include the sites mutated in Cic^6A^. T1059 (blue) was identified as being phosphorylated but was not mutated due to its location in the C2 domain that interacts with ERK.

**Fig. S3.**
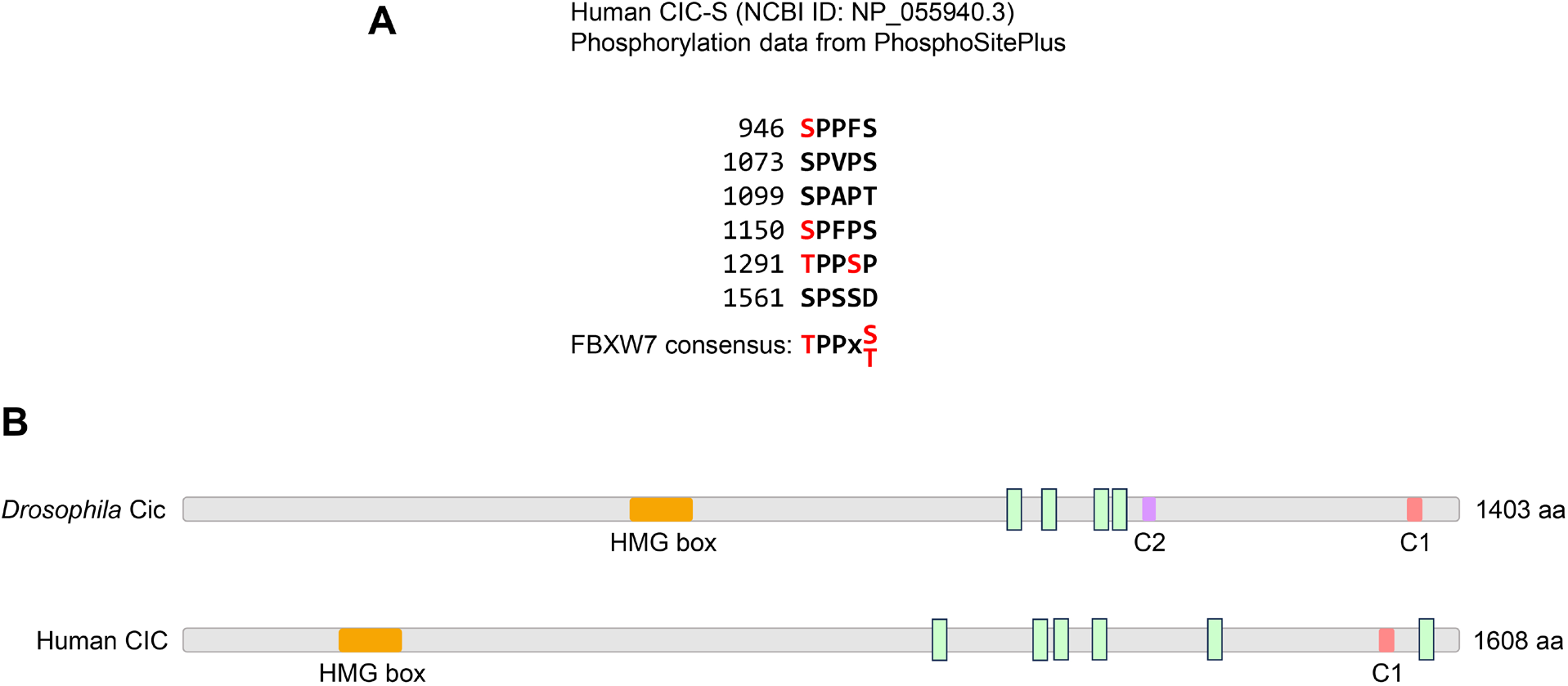
Predicted FBXW7 phosphodegrons in human CIC. (A) Potential FBXW7 phosphodegrons in the human CIC-S protein. Phosphorylated residues are shown in red (data from PhosphoSitePlus). (B) Relative locations of predicted phosphodegrons in *Drosophila* Cic and human CIC proteins. Phosphodegrons are indicated as light green boxes.

